# Study on the effect of a triple cancer treatment of propolis, thermal cycling-hyperthermia, and low-intensity ultrasound on PANC-1 cells

**DOI:** 10.1101/2021.11.19.469215

**Authors:** Yu-Yi Kuo, Wei-Ting Chen, Guan-Bo Lin, Chueh-Hsuan Lu, Chih-Yu Chao

**Author notes:** Correspondence: Chih-Yu Chao Department of Physics, Lab for Medical Physics and Biomedical Engineering, National Taiwan University, No 1, Sec 4, Roosevelt Rd, Taipei 10617, Taiwan, Republic of China (CYC).

## Abstract

**Background:** Pancreatic cancer is a deadly cancer around the world. To reduce side effects and enhance treatment efficacy, study on combination therapy for pancreatic cancer has gained much attention in recent years.

**Methods:** In this paper, we propose a novel triple treatment combining propolis and two physical stimuli‒thermal cycling-hyperthermia (TC-HT) and low-intensity ultrasound (US) on a human pancreatic cancer cell line PANC-1. MTT assay was used to determine the viability of PANC-1 cells. Flow cytometry was used to detect apoptosis, mitochondrial membrane potential (MMP) loss, and intracellular reactive oxygen species (ROS) levels. Western blot analysis was further performed to measure protein expression and phosphorylation.

**Results:** The experiments found that, after the triple treatment, the cell viability of the PANC-1 cells decreased to a level 80% less than the control, without affecting the normal pancreatic cells. Another result was excessive accumulation of ROS after the triple treatment, leading to the amplification of apoptotic pathway through the mitogen-activated protein kinase (MAPK) family and mitochondrial dysfunction. Moreover, the combination of TC-HT and US also promotes the anticancer effect of the heat-sensitive chemotherapy drug cisplatin on PANC-1 cells.

**Conclusion:** This study, to the best of our knowledge, is the first attempt to combine TC-HT, US and a nature compound in cancer treatment. We demonstrate that physical stimuli could augment the therapeutical effect of anticancer agents. It is expected that optimized parameters for different agents and different types of cancer will expand the methodology on oncological therapy in a safe manner.

## Introduction

Cancer is one of the most dreadful diseases and the second leading cause of death around the world. Among all kinds of cancers, pancreatic cancer is the most threatening one, due to its high death rate and low five-year survival rate.^1^ Existing therapies for pancreatic cancer, including surgery, radiation, and chemotherapy, all involve major risks, such as tumor recurrence, refractory, and serious side effects,^2^ as a result of which development of new therapies is of the utmost importance. A popular option is combination therapy, administering two or more anticancer agents to attain a synergistic effect.^3^ However, the interaction between drugs may lead to unexpected competition and even harmful side effects,^4, 5^ jeopardizing patients’ health, let alone improving therapeutic efficacy.

An emerging option in combination therapy is physical stimulus, whose effects on cellular physiology have been reported in several studies.^6–8^ Our team has also looked into the feasibility of combing drug therapy and physical stimuli, such as heat,^9^ electric field,^10^ and magnetic field.^11^ Ultrasound (US) is also a therapeutic tool with extensive application, such as developing internal images,^12, 13^ transporting liposomes to increase agent-delivery rate,^14, 15^ and ablating tumors from normal tissues.^16, 17^ The previous study also found US as a helpful method in inhibiting significantly the viability of PANC-1 cells.^18^ However, US may also entail a number of risks, such as harm to normal tissues around tumors due to overheating caused by high-intensity US.^19, 20^ In addition, liposomes may induce myocardial injury during transport by US.^21, 22^ In view of this, integrating low-intensity US^23, 24^ with a non-hazardous agent is important for expanding the use of US in therapy.

Accordingly, the study employed natural compounds from herbal medicines as anticancer agent. Among natural compounds with therapeutic potential, propolis has been found to be effective in inhibiting several cancer cell lines.^25–27^ In our previous study, propolis was applied, along with thermal cycling-hyperthermia (TC-HT) as a physical stimulus,^28^ in cancer treatment, but the effect still lags behind the in vitro efficacy of chemotherapy drugs.

In this paper, the study introduced low-intensity US as a secondary helper, on top of TC-HT, in order to further augment the anticancer effect of propolis. The novel triple treatment turned out to inhibit the viability of PANC-1 cells significantly, approaching the in vitro efficacy of chemotherapy drugs, without damaging the normal human pancreatic duct cells and skin cells. The study found that the low-intensity US in the triple treatment helped to manipulate the phosphorylation levels of mitogen-activated protein kinase (MAPK) family, thereby activating the intracellular apoptotic signalling. Moreover, it was found in the upstream that intracellular reactive oxygen species (ROS) also increased greatly after the low-intensity US was applied in the triple treatment, thereby boosting the death rate of PANC-1 cells.

## Materials and Methods

### Cell culture and propolis treatment

The human pancreatic cancer cell line PANC-1 and the normal human embryonic skin cell line Detroit 551 were obtained from Bioresource Collection and Research Center (Hsinchu, Taiwan). Normal human pancreatic duct H6c7 cells were purchased from Kerafast, Inc. (Boston, MA, USA). PANC-1 and Detroit 551 cells were cultured respectively in DMEM and EMEM (both from Hyclone, South Logan, UT, USA) supplemented with 10% fetal bovine serum (FBS) (Hyclone) and 1% penicillin-streptomycin (Gibco Life Technologies, Grand Island, NY, USA). H6c7 cells were maintained in keratinocyte-serum free medium (Invitrogen, Life Technologies, Grand Island, NY, USA) supplemented with human recombinant epidermal growth factor, bovine pituitary extract (Invitrogen), and 1% penicillin-streptomycin (Gibco Life Technologies). All cells were maintained in a humidified 5% CO_2_ incubator at 37 °C and subcultured by 0.05% trypsin–0.5 mM EDTA solution (Gibco Life Technologies). Once the confluences reached suitable percentages, cells were plated in 96-well or 35-mm-diameter culture dishes (Thermo Fisher Scientific, Inc., Waltham, MA, USA) for in vitro experiments after 24 h. Propolis was purchased from Grandhealth™ (Grand Health Inc, Richmond, BC, Canada), and cisplatin was obtained from Sigma-Aldrich (St. Louis, MO, USA). All agents were mixed with culture medium to the desired concentration and were incubated with cells for 1h before treating physical stimuli.

### Ultrasound exposure

The US exposure system consisted of a function generator (SG382; Stanford Research Systems, Sunnyvale, CA, USA), a power amplifier (25a250a; Amplifier Research, Souderton, PA, USA), and a planar transducer (A104S-RM; Olympus NDT Inc., Waltham, MA, USA). Continuous pulses were produced using the function generator with the following parameters: -10 dBm amplitude, 1 ms pulse period, and 0.5 ms pulse width. The cell culture plate or dish was placed on the ceramic transducer (resonance frequency 2.25 MHz), which converted electrical signals into acoustic power (**Fig 1A**). To avoid undesirable thermal effects induced by US, the output power of the spatial average intensity of the US exposure was adjusted to be 0.3 W/cm^2^ according to the previous studies.^29, 30^

**Figure 1.**
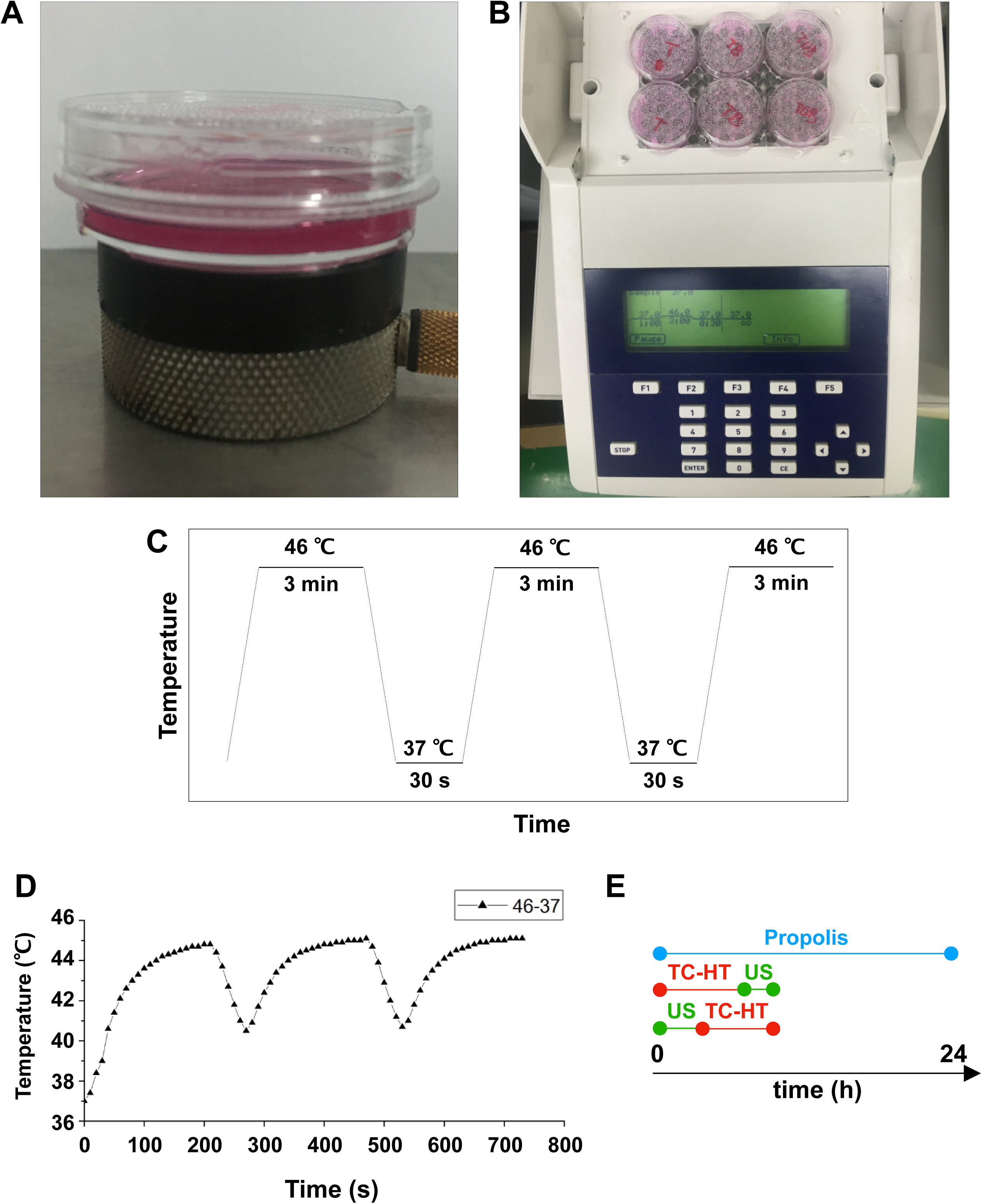
Experimental setups for the triple treatment. **Notes:** 35-mm culture dishes were placed on (**A**) a ceramic transducer and (**B**) a modified PCR machine for the exposures of US and TC-HT, respectively. (**C**) The schematic representation of the TC-HT temperature settings. (**D**) Cell temperature detected by a needle thermocouple when TC-HT was implemented. (**E**) Experimental schedule of the triple treatment with different exposing order of US and TC-HT. **Abbreviations:** US, ultrasound; TC-HT, thermal cycling-hyperthermia.

### Thermal cycling-hyperthermia (TC-HT) treatment

A modified polymerase chain reaction (PCR) system was used to perform TC-HT (Figure 1B). Thermal cycler (model 2720) was purchased from Applied Biosystems (Thermo Fisher Scientific). The system was repeatedly brought to the desired high temperature state and followed by a cooling stage to achieve a series of short period of heat exposure within the desired time (Figure 1C). The experimental setup and administration of TC-HT (10-cycles) have been previously described with optimum results.^28^ The actual temperatures the cancer cells sensed were measured by a needle thermocouple, ranging in 45∼40.5 °C (Figure 1D). Ultrasound exposure was applied either before or after the TC-HT treatment, for different combination tests. During the TC-HT treatment (∼ 45 min), the control and propolis-treated groups were under room temperature (RT) without a 5% CO_2_ environment. After the treatments, cells were maintained in the cell culture incubator for the following experiments.

### MTT assay

3-(4,5-dimethylthiazol-2-yl)-2,5-diphenyltetrazolium bromide (MTT) (Sigma-Aldrich) was dissolved in distilled water to prepare a 5 mg/ml stock solution. The treated PANC-1 cells were incubated for 4 h at 37 °C with a final MTT concentration 0.5 mg/ml in DMEM culture medium to assess the cell viabilities. The formazan crystals were dissolved by equal volume of the solubilizing buffer of 10% sodium dodecyl sulfate (SDS) (Bioshop Canada Inc., Burlington, ON, Canada) solution in 0.01 N hydrochloric acid (HCl) (Echo Chemical Co. Ltd., Miaoli, Taiwan) at 37 °C overnight. The absorbance of each well was detected by Multiskan GO microplate Spectrophotometer (Thermo Fisher Scientific), and the quantity of formazan was determined by the absorbance at 570 nm, with a background subtraction at 690 nm. The cell viabilities were expressed in percentage and the untreated control was set at 100%.

### Treatment with ROS scavenger

PANC-1 cells were seeded into 96-well or 35-mm-diameter culture dishes overnight. For ROS inhibition analysis, cells were pretreated with 5 mM N-acetyl-cysteine (NAC) (Sigma-Aldrich) in culture medium for 1 h and subsequently treated with propolis and/or physical stimuli.

### Apoptotic analysis by flow cytometry

PANC-1 cells were collected 24 h after treatments and then rinsed twice with ice-cold phosphate buffered saline (PBS) (Hyclone). The apoptotic rates were analyzed by the Annexin V-FITC and propidium iodide (PI) double detection kit (BD Biosciences, San Jose, CA, USA), and the rinsed cells were resuspended in binding buffer containing Annexin V-FITC and PI and then incubated at RT for 15 min in the dark. Apoptotic signals were detected by FACSCanto II system (BD Biosciences).

### ROS and mitochondrial membrane potential (MMP) analyses by flow cytometry

ROS was detected using the fluorescent dye dihydroethidium (DHE) (Sigma-Aldrich), and the loss of MMP was determined using the lipophilic cationic fluorescent dye 3,3’-dihexyloxacarbocyanine iodide (DiOC_6_(3)) (Enzo Life Sciences, Inc., Plymouth Meeting, PA, USA). PANC-1 cells were harvested 24 h after treatments and rinsed with PBS before staining. Rinsed cells were resuspended and then incubated with 5 μM DHE or 20 nM DiOC_6_(3) in PBS at 37 °C for 30 min in the dark. The fluorescence signals were measured by FACSCanto II system (BD Biosciences) with the PE channel (for DHE staining) or FL1 channel (for DiOC_6_(3) staining).

### Western blot analysis

Protein expression levels of PANC-1 cells were quantified by western blot analysis. Cells were rinsed with PBS and then lysed in the lysis buffer (50 mM Tris-HCL, pH 7.4, 0.15 M NaCl, 0.25% deoxycholic acid, 1% NP-40, 1% Triton X-100, 0.1 % SDS, 1 mM EDTA) (Millipore, Billerica, MA, USA), supplemented with active protease (Millipore) and phosphatase inhibitor cocktail (Cell signaling Technology, Danvers, MA, USA). After centrifugation, the supernatants were collected and the protein concentrations were quantified by Bradford protein assay (Bioshop, Inc.). Equal amount of proteins (20 μg) were resolved by 10% SDS-polyacrylamide gel electrophoresis (SDS-PAGE) and then transferred onto polyvinylidene fluoride (PVDF) membranes (Millipore). 5% skim milk powder or 5% bovine serum albumin in TBST (20 mM Tris-base, pH 7.6, 0.15 M NaCl, 0.1% Tween 20) was used to block nonspecific antibody binding sites for 1 h at RT. Afterwards, the blocked membranes were probed with specific primary antibodies against phosphorylated extracellular signal-regulated kinases (p-ERK), phosphorylated c-Jun N-terminal kinase (p-JNK), poly (ADP-ribose) polymerase (PARP) (Cell signaling), phosphorylated p38 MAPK (p-p38), and glyceraldehyde-3-phosphate dehydrogenase (GAPDH) (Gentex, Irvine, CA, USA) at 4 °C overnight. The membranes were rinsed with TBST buffer three times and then incubated with horseradish peroxidase-conjugated goat anti-rabbit secondary antibodies (Jackson ImmunoResearch Laboratories, West Grove, PA, USA) in a blocking solution at RT for 1 h. Immunoreactivity signal was amplified by an enhanced chemiluminescence (ECL) substrate (Advansta, San Jose, CA, USA) and detected by an imaging system Amersham Imager 600 (GE Healthcare Life Sciences). GAPDH was used as the loading control to normalize the relative folds of targeting proteins.

### Statistical analysis

Experiments were repeated three times for validation, and statistical analyses were performed using one-way analysis of variance (ANOVA) by OriginPro 2015 software (OriginLab). Results were expressed as the mean ± standard deviation, and were considered to be statistically significant when *p*-values were less than 0.05.

## Results

### Triple treatment greatly inhibits the viability of PANC-1 cells

The cell viability of PANC-1 cells versus the propolis concentration was performed in a gradient manner. When the propolis was less than 0.5%, as shown in Figure 2A, there was no notable inhibition effect on MTT results. However, when it exceeded 0.5%, the cell viability dropped significantly. Therefore, a moderate propolis concentration 0.3% was chosen for the following experiments. Next, the physical stimuli of TC-HT and low-intensity US were introduced to affect the viability of PANC-1 cells. In our study, 10-cycles TC-HT and 2.25 MHz US with intensity 0.3W/cm^2^ and duration 30 minutes were chosen to avoid the thermotoxicity on PANC-1 cells. As shown in Figure 2B, we found that the combination of TC-HT and US was also innocent to PANC-1 cells, but when 0.3% propolis was involved in the triple treatment, the viability of PANC-1 cells was greatly inhibited. It was also noted that the implementation order of TC-HT and US in triple treatment was influential. When TC-HT was performed prior to US (TC-HT + US) in the presence of 0.3% propolis, US helped to further suppress the cell viability of PANC-1 cells significantly down to 17.1%, cutting more than 80% of the viability of the untreated control and thus approaching the in vitro efficacy of chemotherapy drugs. In comparison, the treatment that US was performed prior to TC-HT (US + TC-HT) showed a less inhibition effect (43.1% viability), and hence we adopted the implementation order TC-HT + US as the protocol of the triple treatment in the subsequent experiments. Furthermore, in all double treatments, only 0.3% propolis + TC-HT showed notable inhibition effect (48.9% viability) on PANC-1 cells, which was consistent with our previous results.^28^ However, 0.3% propolis + US performed a relatively poor inhibition effect (65.4% viability) on PANC-1 cells, and as a result it was also not included in the following experiments. Figure 2C showed the light microscope images of PANC-1 cells 24 h after each treatment, and the cell morphologies demonstrated an evident inhibition effect on PANC-1 cells after the triple treatment. Moreover, normal cells such as the human skin cells Detroit 551 (Figure 2D) and human pancreatic duct cells H6c7 (Figure 2E) were not significantly affected by the triple treatment as well as all the other treatments. The result indicates that the triple treatment could have a good selective effect on carcinoma cells and normal cells, which makes it safer and more feasible in anticancer treatment.

**Figure 2.**
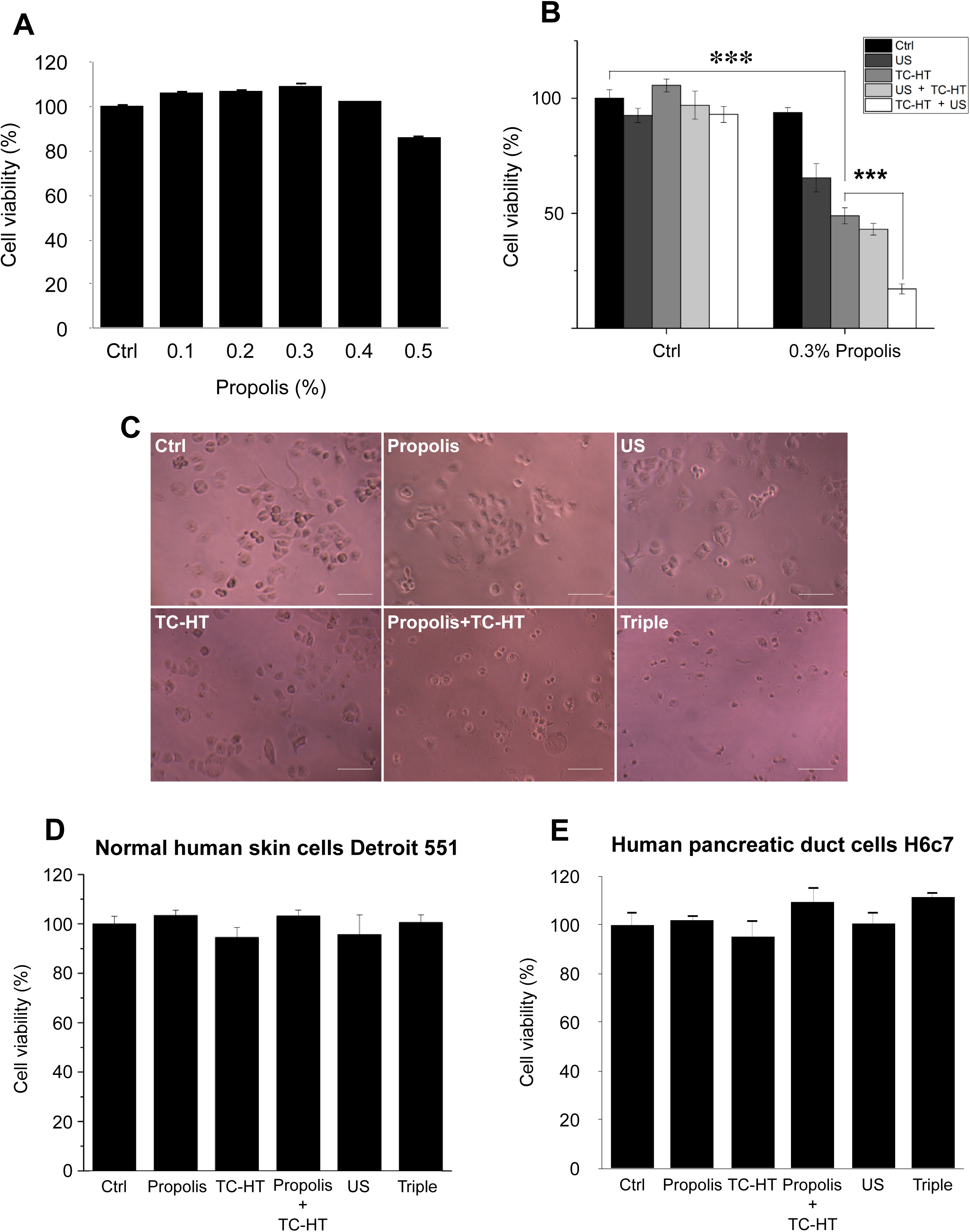
Viability inhibition effects of propolis on PANC-1, Detroit 551, and H6c7 cells. **Notes:** MTT assay was conducted to determine the viabilities of PANC-1 cells after the treatment of (**A**) different propolis concentrations and (**B**) different combinations of physical stimulations. (**C**) Representative light microscope images of PANC-1 cells after each treatment, scale bar = 100 μm. The viabilities of (**D**) normal human skin cells Detroit 551 and (**E**) normal human pancreatic duct cells H6c7 were measured 24 h after each treatment. Data were presented as the mean ± standard deviation in triplicate (***P<0.001). Ctrl, Control group with no treatment; Propolis, Propolis group with 0.3% (v/v) propolis; TC-HT, TC-HT group with 10 cycles TC-HT (setting: 46 °C 3 min – 37 °C 30 s); Propolis + TC-HT, Propolis + TC-HT group with 0.3% (v/v) propolis and 10 cycles TC-HT (setting: 46 °C 3 min – 37 °C 30 s), sequentially; US, US group with 2.25 MHz US (0.3 W/cm^2^) for 30 min; Triple, Triple group with 0.3% (v/v) propolis, 10 cycles TC-HT (setting: 46 °C 3 min – 37 °C 30 s), and 2.25 MHz US (0.3 W/cm^2^) for 30 min, sequentially. **Abbreviations:** US, ultrasound; TC-HT, thermal cycling-hyperthermia.

### Triple treatment increases intracellular ROS levels synergistically

Intracellular ROS is an important regulator of cell death. It has been reported that heat and low-intensity US could elevate the intracellular ROS level.^31, 32^ We further investigated whether ROS was increased in response to the triple treatment, so the DHE was used in this experiment to determine the level of superoxide radical anion (O_2_^‧-^) in PANC-1 cells after each treatment. As shown in Figure 3A-B, it was found that propolis hardly changed the fluorescence signals. Although propolis + TC-HT significantly deformed the fluorescence intensity distribution in an enhanced manner (1.6-fold to control), it did not significantly differ from the enhancement induced by TC-HT alone. In addition, US elevated ROS levels as well, though not as many as TC-HT. Noticeably, the triple treatment showed a significant accumulation of the intracellular ROS (up to 2.1-fold of the control group), which was also significantly higher than the TC-HT + 0.3% propolis treatment. The result suggested that, in the triple treatment, US helped to further boost up the generation of ROS in PANC-1 cells, and could result in enhanced cell death rate after the treatment.

**Figure 3.**
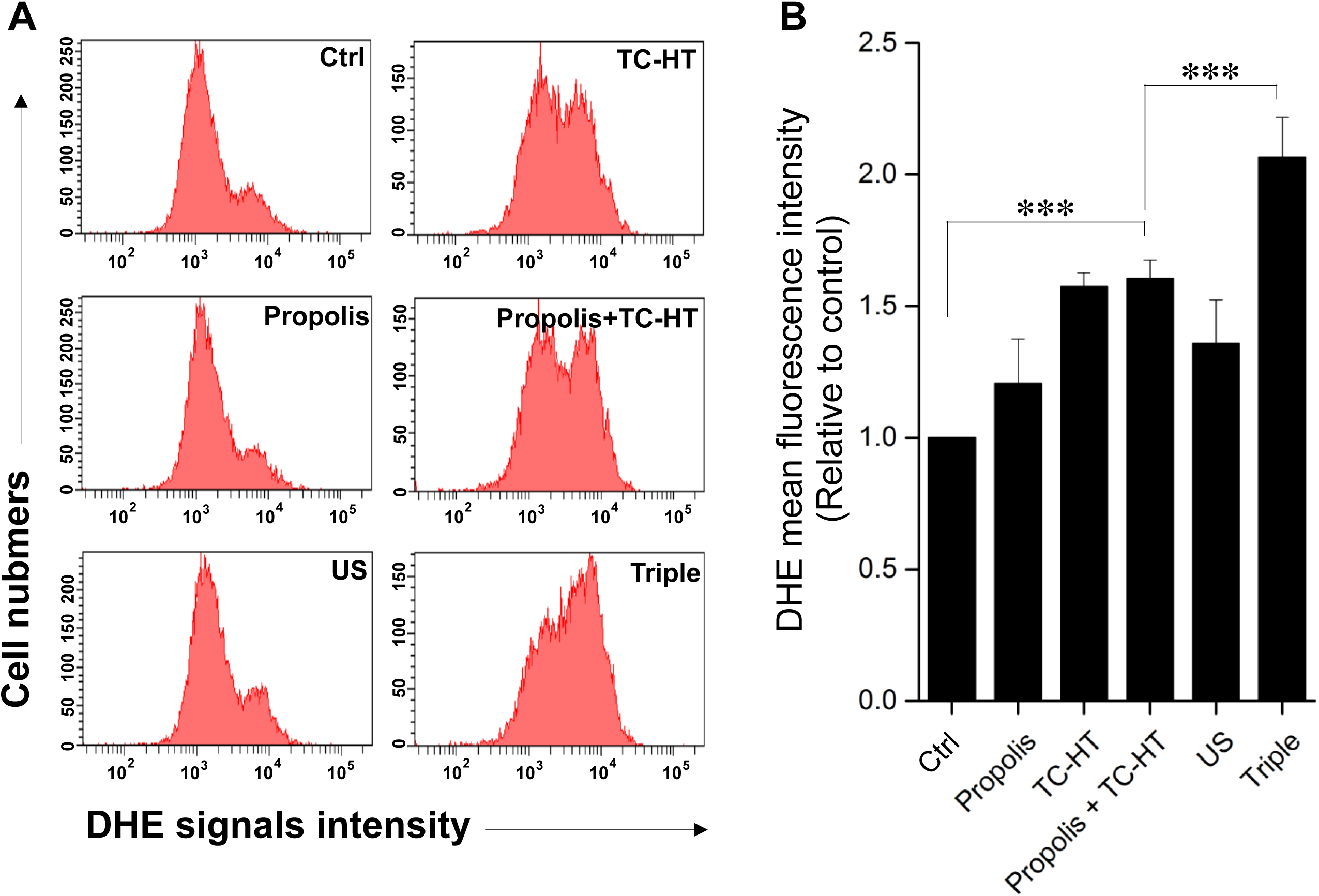
The triple treatment raised the generation of intracellular ROS. **Notes:** (**A**) Intracellular superoxide radical anion (O_2_^‧-^) levels of PANC-1 cells were determined by flow cytometry with the fluorescent dye DHE. (**B**) Quantification of the mean DHE fluorescence levels after each treatment. Data were presented as the mean ± standard deviation in triplicate (***P<0.001). Ctrl, Control group with no treatment; Propolis, Propolis group with 0.3% (v/v) propolis; TC-HT, TC-HT group with 10 cycles TC-HT (setting: 46 °C 3 min – 37 °C 30 s); Propolis + TC-HT, Propolis + TC-HT group with 0.3% (v/v) propolis and 10 cycles TC-HT (setting: 46 °C 3 min – 37 °C 30 s), sequentially; US, US group with 2.25 MHz US (0.3 W/cm^2^) for 30 min; Triple, Triple group with 0.3% (v/v) propolis, 10 cycles TC-HT (setting: 46 °C 3 min – 37 °C 30 s), and 2.25 MHz US (0.3 W/cm^2^) for 30 min, sequentially. **Abbreviations:** US, ultrasound; TC-HT, thermal cycling-hyperthermia; DHE, dihydroethidium.

### Triple treatment increases mitochondrial apoptosis in PANC-1 cells

It has been known that the enhanced intracellular ROS levels were positively correlated to mitochondrial apoptosis.^33^ In our work, the apoptotic rates of PANC-1 cells after various treatments were analyzed by the flow cytometry with the fluorescence dye Annexin V and PI (Figure 4A-B). With the aid of US, the triple treatment further caused 55.3% apoptotic rate, which was significantly higher than the 23.5% apoptotic rate caused by the double treatment of propolis and TC-HT. The cell apoptosis results observed here were highly consistent with the results of the accumulated ROS levels in PANC-1 cells after the same treatment, as described in Figure 3. Furthermore, the mitochondrial membrane potential (MMP) was assessed using DiOC_6_(3) fluorescence staining by flow cytometric analysis. As shown in Figure 4C-D, the ratio of the cells exhibiting MMP loss was significantly promoted to 23.3% after the double treatment of propolis + TC-HT, and it was further elevated significantly to 34.7% by employing the triple treatment. These results showed that adopting US in the triple treatment could decrease MMP level, and hence caused more mitochondrial dysfunction. The decreased MMP level was an indicator of mitochondrial apoptosis, and since the results of apoptosis assay (Figure 4A-B) and MMP assay (Figure 4C-D) were quite similar, we believe that the mitochondrial dysfunction was implicated in the apoptosis of PANC-1 cells via the triple treatment.

**Figure 4.**
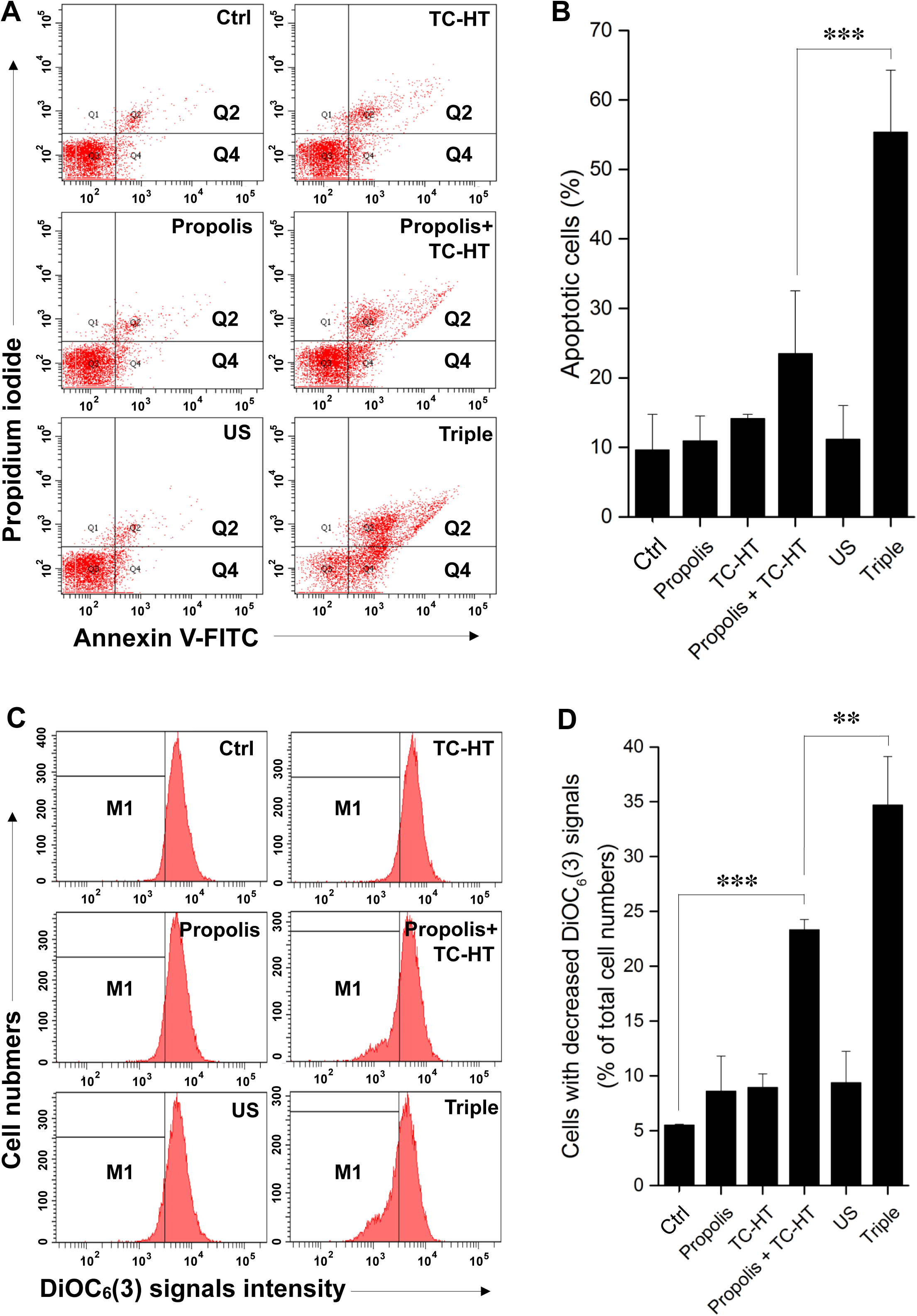
Triple treatment increased apoptosis on PANC-1 cells via mitochondrial dysfunction. **Notes:** (**A**) Apoptosis after treatment was analyzed via flow cytometry with Annexin V-FITC/PI double staining, and (**B**) the apoptotic percentage (Q2+Q4) were calculated. (**C**) MMP level after treatment was analyzed via flow cytometry with DiOC_6_(3) staining, and (**D**) the percentage of cells with the loss of MMP (M1) was calculated. Data were presented as the mean ± standard deviation in triplicate (***P<0.001, **P<0.01). Ctrl, Control group with no treatment; Propolis, Propolis group with 0.3% (v/v) propolis; TC-HT, TC-HT group with 10 cycles TC-HT (setting: 46 °C 3 min – 37 °C 30 s); Propolis + TC-HT, Propolis + TC-HT group with 0.3% (v/v) propolis and 10 cycles TC-HT (setting: 46 °C 3 min – 37 °C 30 s), sequentially; US, US group with 2.25 MHz US (0.3 W/cm^2^) for 30 min; Triple, Triple group with 0.3% (v/v) propolis, 10 cycles TC-HT (setting: 46 °C 3 min – 37 °C 30 s), and 2.25 MHz US (0.3 W/cm^2^) for 30 min, sequentially. **Abbreviations:** US, ultrasound; TC-HT, thermal cycling-hyperthermia; DiOC_6_(3), 3,3’-dihexyloxacarbocyanine iodide.

### The apoptosis induced by triple treatment is regulated through MAPK pathway

The activation of apoptotic signalling was examined by western blot analysis. As shown in Figure 5A, we found that the PARP cleavage was significantly increased (2.9-fold of control) after propolis + TC-HT treatment on PANC-1 cells. Noticeably, the PARP cleavage was further promoted significantly to 6.2-fold of control by US in the triple treatment (Figure 5A). Together with the previous flow cytometry results of apoptosis and MMP, it was pointed out that propolis + TC-HT could activate the mitochondrial apoptosis signalling in PANC-1 cells, and US in the triple treatment could further help this cascade to realize a near-chemotherapy level treatment in vitro.

**Figure 5.**
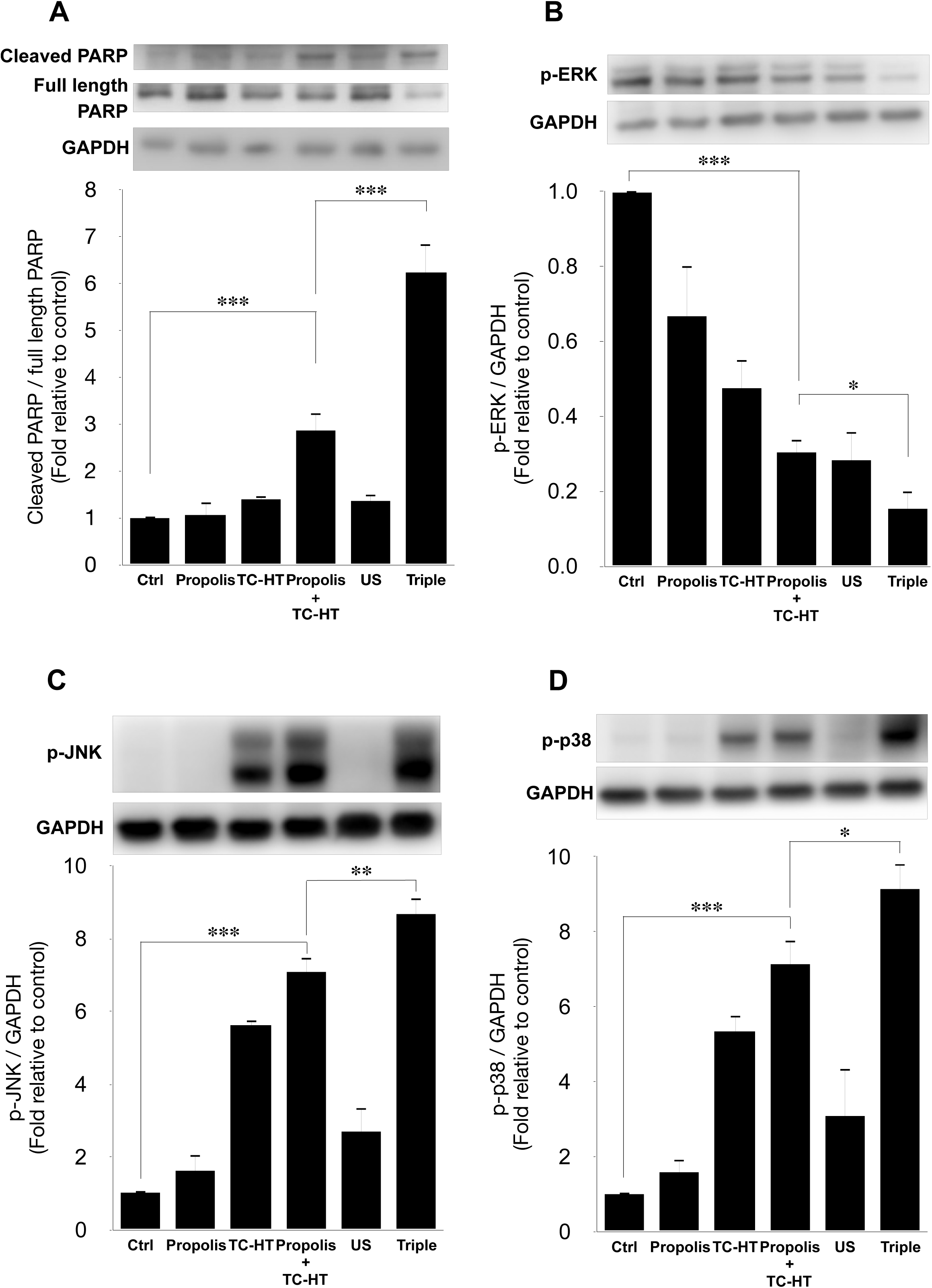
Triple treatment modulated apoptosis via regulating the MAPK family. **Notes:** Representative western blots of the apoptosis-related proteins and the quantification of (**A**) the PARP cleavage (ratio of cleaved PARP/full length PARP), the phosphorylation level of (**B**) ERK, (**C**) JNK, and (**D**) p38. GAPDH was used as loading control. Data were presented as the mean ± standard deviation in triplicate (***P<0.001, **P<0.01, *P<0.05). Ctrl, Control group with no treatment; Propolis, Propolis group with 0.3% (v/v) propolis; TC-HT, TC-HT group with 10 cycles TC-HT (setting: 46 °C 3 min – 37 °C 30 s); Propolis + TC-HT, Propolis + TC-HT group with 0.3% (v/v) propolis and 10 cycles TC-HT (setting: 46 °C 3 min – 37 °C 30 s), sequentially; US, US group with 2.25 MHz US (0.3 W/cm^2^) for 30 min; Triple, Triple group with 0.3% (v/v) propolis, 10 cycles TC-HT (setting: 46 °C 3 min – 37 °C 30 s), and 2.25 MHz US (0.3 W/cm^2^) for 30 min, sequentially. **Abbreviations:** US, ultrasound; TC-HT, thermal cycling-hyperthermia.

Moreover, it was known that the PARP cleavage could be modulated by MMP level, and mitochondrial dysfunction could also be regulated by the excessive intracellular ROS via MAPK family.^34^ In MAPK family, the p-ERK level represented the activation of cell survival,^35^ while the p-JNK and p-p38 levels were the indicator of cell death.^36, 37^ In this study, it was found that the p-ERK level was suppressed by propolis + TC-HT treatment (0.30-fold), and was further down-regulated when US was introduced in the triple treatment (0.15-fold) (Figure 5B). In addition, the p-JNK and p-p38 levels both exhibited a reverse performance, which were promoted the most in the triple treatment (8.7-fold & 9.2-fold, respectively) (Figure 5C-D). These results were consistent with the results of ROS and MMP assessments by flow cytometry. Therefore, we speculated that the excess intracellular ROS induced by the triple treatment regulated the activation of the MAPK family and thus caused mitochondrial dysfunction and the cascade of apoptosis.

### ROS scavenger attenuates the apoptosis induced by triple treatment

To further confirm that the cell death after the triple treatment was regulated by the generation of intracellular ROS, the ROS scavenger NAC was applied in the experiment.^38^ 5 mM NAC was incubated with PANC-1 cells 1 h prior to the triple treatment. As shown in Figure 6A, the inhibitory effect of the triple treatment was restored by NAC, and NAC itself did not affect the viability of PANC-1 cells. Similar results were also observed in the activation of apoptotic pathway, as shown in Figure 6B. NAC alone did not affect PARP cleavage, but it significantly down-regulated the triple treatment-promoted PARP cleavage (Figure 6B). Therefore, the results supported our speculation that the triple treatment could induce mitochondrial apoptosis of PANC-1 cells via the excessive increment of intracellular ROS.

**Figure 6.**
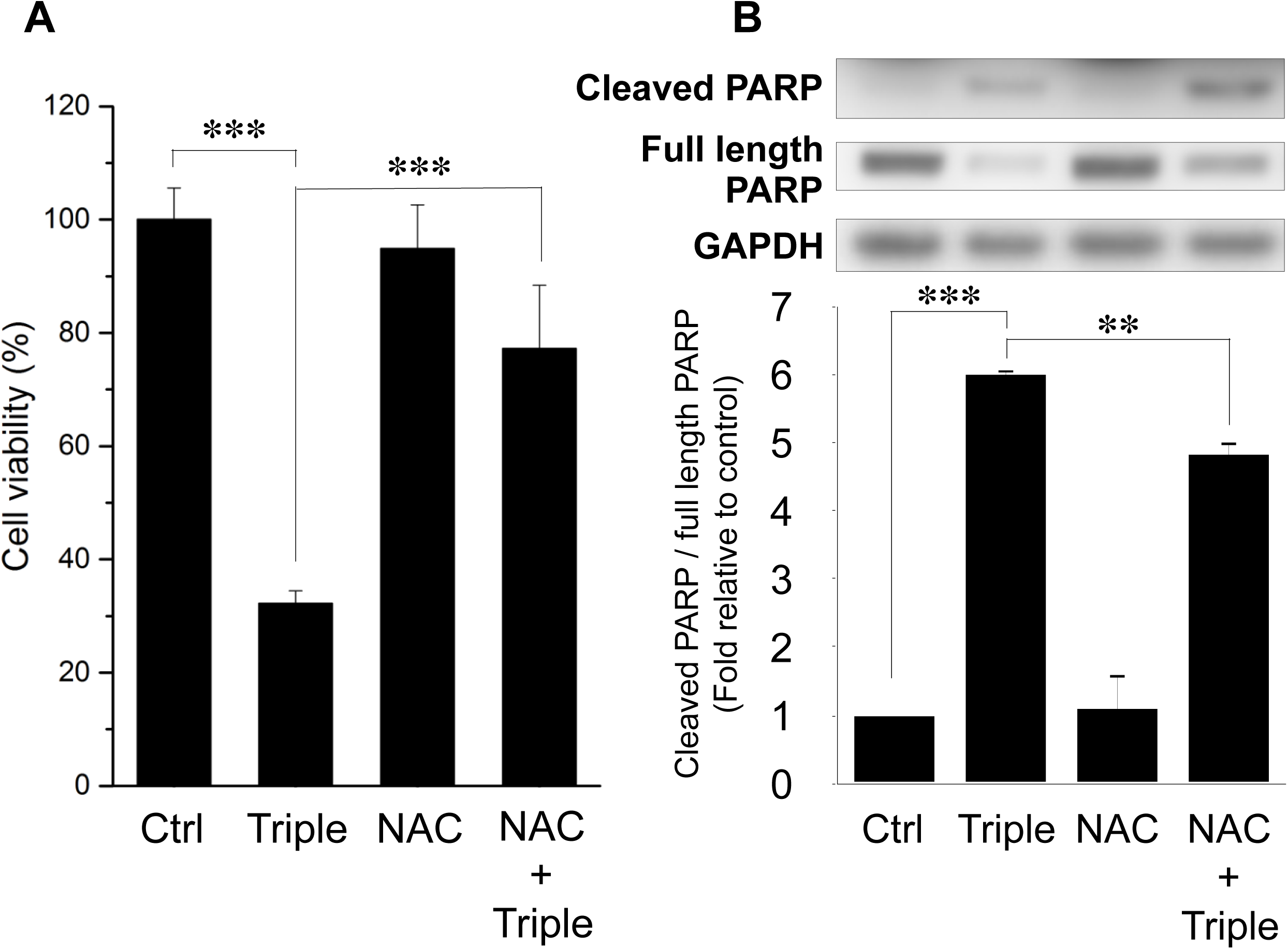
ROS inhibition blocked the cell death and the activation of apoptosis pathways. **Notes:** (**A**) Cell viabilities of PANC-1 cells and (**B**) the quantification of PARP cleavage (ratio of cleaved PARP/full length PARP) after the triple treatment with or without 1h NAC pretreatment. GAPDH was used as loading control. Data were presented as the mean ± standard deviation in triplicate (***P<0.001, **P<0.01). Ctrl, Control group with no treatment; Triple, Triple group with 0.3% (v/v) propolis, 10 cycles TC-HT (setting: 46 °C 3 min – 37 °C 30 s), and 2.25 MHz US (0.3 W/cm^2^) for 30 min, sequentially. NAC, NAC group with 5 mM NAC; NAC + Triple, NAC + Triple group with 5 mM NAC prior to the triple treatment. **Abbreviation:** NAC, N-acetyl-cysteine.

### Triple treatment can be applied with chemotherapy drug as a novel anticancer treatment

In this study, we have shown that the method TC-HT followed by mild US exposure could further amplify the anticancer effect of propolis. But, the question is whether the TC-HT + US method can be expanded to the existing chemotherapy drugs. Cisplatin, for instance, was a commonly used clinical chemotherapy drug for several kinds of cancers such as lung, ovarian, breast, and brain cancer.^39^ Besides, it has also been reported that cisplatin was sensitive to heat.^40^ However, the conventional therapeutic dosage of cisplatin could cause severe side effects to the patients,^41, 42^ and therefore it was important to develop a new method to reduce the effective dose of cisplatin. As a result, we applied the method of TC-HT + US with cisplatin on the PANC-1 cells to investigate the potential of this triple treatment method. A relatively low dose of 1 μM cisplatin and short incubation time 24 h was chosen for the MTT assay independently or in combination with the physical stimulations. ^43, 44^ After the treatment, the viability of PANC-1 cells was just slightly inhibited by 7.5% by individual cisplatin, and the double treatment of cisplatin + TC-HT also did not alter the viability of PANC-1 cells. The triple treatment, however, promoted the inhibitory effect significantly up to 48.2% (Figure 7). Compared to the conventional results of cisplatin, our method could not only reduce the effective dose but also boost up the anticancer effect of cisplatin. Therefore, the method of physical stimuli in the study should hold great potential to apply onto other heat sensitive chemotherapy drugs or anticancer agents to achieve a better anticancer effect.

**Figure 7.**
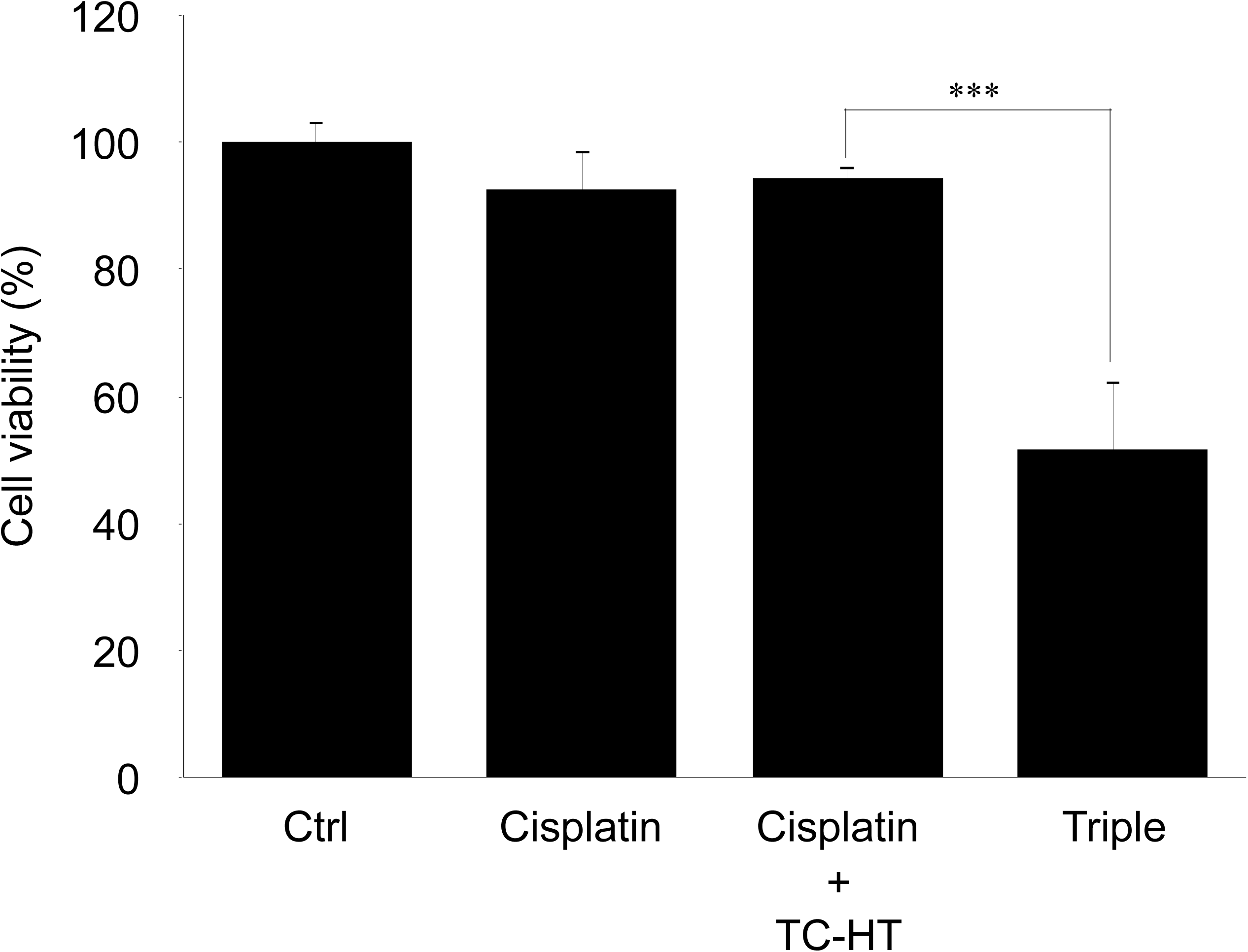
The method of triple treatment promoted the inhibitory effect of the heat sensitive chemotherapy drug cisplatin. **Notes:** Cell viabilities of PANC-1 cells treated with cisplatin, cisplatin + TC-HT, and the triple treatment of cisplatin + TC-HT + US. Cisplatin concentrations in all treatments were 1 μM. Data were presented as the mean ± standard deviation in triplicate (***P<0.001). Ctrl, Control group with no treatment; Cisplatin, Cisplatin group with 1 µM cisplatin; Cisplatin + TC-HT, Cisplatin + TC-HT group with 1 µM cisplatin, 10 cycles TC-HT (setting: 46 °C 3 min – 37 °C 30 s), sequentially; Triple, Triple group with 1 µM cisplatin, 10 cycles TC-HT (setting: 46 °C 3 min – 37 °C 30 s), and 2.25 MHz US (0.3 W/cm^2^) for 30 min, sequentially. **Abbreviation:** TC-HT, thermal cycling-hyperthermia.

## Discussion

Combination treatment augments the anticancer effect of individual agents by activating multiple pathways, thereby lowering the necessary dosage of the agents to a level harmless to normal cells and human health. However, research on combination anticancer agents is costly and time-consuming.^45^ Moreover, unpredictable molecular interactions may be detrimental to patients’ health.^5^ Therefore, the application of physical stimuli has been considered as a potential candidate for combinative anticancer treatments in combating cancer. Following several combination treatments of physical stimuli and herbal compounds proposed by our team previously,^46^ the study put forth the combination treatment of propolis, TC-HT, and low-intensity US, proving that it can inhibit pancreatic cancer cell line PANC-1 at a level close to the in vitro efficacy of chemotherapy drugs.

In cancer cells, intracellular ROS levels have been known to be the main source of the oxidative stress,^47^ a key factor for cell viability.^48^ Elevated ROS level has been shown to be able to activate some signalling pathways associated with cell proliferation, apoptosis, and cell cycle progression.^49^ Besides, heat treatment and low-intensity US both can increase the intracellular ROS levels.^31, 32^ In this work, the study shows that TC-HT significantly augments the intracellular ROS levels of PANC-1 cells (Figure 3A-B), at an extent higher than the individual effects of propolis and low-intensity US. It was found that the combination of propolis and TC-HT did not further elevate the levels in PANC-1 cells. Nevertheless, after the low-intensity US was administered, the triple treatment showed a great improvement effect, doubling the intracellular ROS levels of PANC-1 cells. It has been known that excessive intracellular ROS level could activate the apoptotic pathway cascade, increasing the apoptosis in the carcinoma cells.^33^ Our study demonstrated that the apoptotic rate of PANC-1 cells was elevated along with the increase of the intracellular ROS levels. The result showed that the triple treatment induced the highest apoptotic rate, compared with other approaches, suggesting its ability to regulate the death of PANC-1 cells via excessive intracellular ROS accumulation. To demonstrate the crucial role of the intracellular ROS levels, NAC was employed in the following experiments on the triple treatment. While independent NAC pretreatment did not affect the viabilities of PANC-1 cells (Figure 6A), it protected the cells from cytotoxicity in the triple treatment. Hence, ROS elevation played a key role in the anticancer effect in the triple treatment.

The initiation of apoptosis, a common cell death mechanism, is closely related to the function of mitochondria, which is the chemical-energy source of cells and critical for the viability of cells.^50^ It was reported that US could induce mitochondrial dysfunction,^51^ and the dysfunction could induce a series of biochemical cascade of apoptosis, thereby blocking cell proliferation.^52^ Meanwhile, the activated members of the MAPK family, such as ERK, JNK, and p38 were deemed to be capable of regulating the dysfunction of mitochondria, and the elevated ROS level was shown to be conducive to the activation of p38 and JNK but down-regulate the activation of ERK.^53^ The results suggest that excessive ROS further induced by US could cause greater mitochondrial apoptotic rate via additionally activating the MAPK family members. In this work, our study showed that adopting US in the triple treatment raised greater apoptotic rate of PANC-1 cells (Figure 4A-B), while decreasing the MMP level lower, which led to more severe dysfunction of mitochondria than the double treatment of propolis and TC-HT (Figure 4C-D). Furthermore, the mild US in the triple treatment further helped to increase the phosphorylated levels of p38 and JNK significantly, while inhibiting the phosphorylation of ERK (Figure 5), underscoring its ability to manipulate the function of mitochondria via the ROS-activated MAPK family proteins.

The injured mitochondria would release cytochrome-c into the cytoplasm,^54^ cleaving caspase 9 and thus activating caspase 3 in the downstream,^55^ which entered further the nucleus and cleaved PARP. Then PARP would lose its enzyme activity, initiating apoptosis irreversibly.^56^ In addition, it was reported that US could induce mitochondrial apoptosis in cancer cells.^51, 57^ The study demonstrated that the triple treatment could induce mitochondrial dysfunction via regulating the phosphorylation level of the members of MAPK family. In addition, the triple treatment also increased PARP cleavage significantly, compared with the combination treatment of propolis and TC-HT (Figure 5A). Furthermore, as shown in Figure 6B, it was found that the triple treatment-promoted PARP cleavage was significantly suppressed by NAC, which indicates that increased ROS level was the key regulator for the apoptotic effect of the triple treatment.

The triple treatment in the study included two physical stimuli, TC-HT and US, which could be integrated for simultaneous implementation via the high-intensity focused ultrasound (HIFU). Being able to raise temperature in the exposed region for energy transfer,^19, 20^ HIFU has been applied for tumor ablation for over a decade.^16, 58^ The temperature increase could be electrically controlled and directed to the targeted region.^59^ Therefore, with the help of HIFU, the triple treatment could be applied clinically. To augment the effect, cisplatin, a common thermal sensitive chemotherapy drug, was incorporated as a substitute for propolis into the triple treatment. It was shown that the inhibited viability of PANC-1 cells by cisplatin was further suppressed with a large extent when both of TC-HT and US were introduced into the treatment (Figure 7). The dosage of cisplatin can be reduced to a lower concentration, without compromising the anticancer effect. As a result, the triple treatment has the potential supplementing the administration of drugs, not only augmenting the effect but also reducing the dosage of the latter.

## Conclusion

In summary, this study proposed for the first time an effective triple cancer treatment combining propolis, TC-HT, and low-intensity US, which could significantly suppress the growth of PANC-1 cells via an ROS-modulated mitochondrial apoptosis, with a performance comparable to chemotherapy. The study also showed that the triple treatment could induce mitochondrial dysfunction via the regulation of MAPK family, resulting in apoptosis via the up-regulated PARP cleavage. It also demonstrated that the ROS level plays a key role in the performance of the triple treatment. In addition, chemotherapy drugs, such as cisplatin, can be incorporated into the treatment as substitute for propolis. The triple treatment incorporating cisplatin also exhibited a much higher effect in inhibiting cancer cell growth than the cisplatin alone, promising to increase the performance and safety of the existing cancer therapy. Overall, the study proposed employment of physical stimuli, as a promising option in cancer therapy.

## Acknowledgments

The authors would like to acknowledge the service provided by Technology Commons in College of Life Science, National Taiwan University for use of flow cytometry system. This work was supported by grants from Ministry of Science and Technology (MOST 110-2112-M-002-004, MOST 109-2112-M-002-004, and MOST 108-2112-M-002-016 to CYC) and Ministry of Education (MOE 106R880708 to CYC) of the Republic of China. The funders had no role in study design, data collection and analysis, decision to publish, or preparation of the manuscript.

